# Cell-type-specific expression of classical and membrane progesterone receptors at the human maternal–fetal interface

**DOI:** 10.64898/2026.09.21.752943

**Authors:** Akihito Sagara, Risa Shimokawa, Yutaka Iwagoi, Masaru Kobayashi, Munekage Yamaguchi, Haruta Mogami, Fumitaka Osakada, Kentaro Nagaoka, Hiroaki Ohno, Ikuo Kimura, Eiji Kondoh

## Abstract

Progesterone is essential for the establishment and maintenance of pregnancy, primarily through the classical nuclear progesterone receptor (PGR). However, progesterone also exerts rapid, non-genomic effects through membrane progesterone receptors, including members of the progestin and adipoQ receptor (PAQR) family, whose cellular distribution and potential roles at the human maternal–fetal interface remain poorly understood. We therefore sought to define the cell-type-specific expression landscape of classical and membrane progesterone receptors during early human pregnancy.

We analyzed publicly available single-cell RNA-sequencing datasets of the first-trimester human maternal–fetal interface to characterize the expression of PGR and PAQR family members across trophoblast, immune, stromal, and vascular cell populations. Major expression patterns were examined across independent datasets and further assessed using spatial transcriptomic data.

PGR expression was predominantly localized to decidual stromal and perivascular populations, whereas individual PAQR family members showed distinct cellular distributions. PAQR6 was enriched in decidual natural killer cells, PAQR7 was broadly expressed in trophoblast populations, PAQR8 was prominently expressed in fetal stromal and macrophage populations, including Hofbauer cells, and PAQR9 showed prominent enrichment in syncytiotrophoblasts. Selected cell-type-specific patterns were further supported by spatial transcriptomic analysis.

These findings provide a cell-type-resolved framework for investigating how distinct progesterone receptor systems mediate progesterone signaling across maternal and fetal cellular compartments during early human placentation.

## Introduction

Progesterone plays essential roles in the establishment and maintenance of pregnancy, including endometrial decidualization, modulation of the maternal immune environment, and placental development [1,2]. These actions have been studied predominantly in the context of transcriptional regulation mediated by the classical nuclear progesterone receptor (PGR) [1,3]. However, progesterone also exerts rapid, non-genomic actions through membrane-associated signaling mechanisms. Among the receptors implicated in these actions are five members of the progestin and adipoQ receptor (PAQR) family, collectively referred to as membrane progestin receptors (mPRs): PAQR5–PAQR9 [4]. Thus, progesterone actions in gestational tissues may involve multiple receptor systems, including both classical PGR and membrane progestin receptors.

During the first trimester, the human maternal–fetal interface comprises diverse fetal and maternal cell populations, including trophoblasts, fetal stromal cells, Hofbauer cells, decidual stromal cells, decidual natural killer (dNK) cells, and macrophages. These cell populations interact to support placental development, trophoblast invasion, and establishment of the local immune environment [5,6]. Single-cell RNA sequencing studies by Vento-Tormo et al. and Suryawanshi et al. have provided detailed transcriptomic atlases of the first-trimester placenta and decidua, enabling the maternal–fetal interface to be examined at cellular resolution [5,6]. Although individual membrane progestin receptors have previously been detected in endometrial and gestational tissues [7], the cell-type-specific distribution of classical PGR and PAQR family members across the human first-trimester maternal–fetal interface has not been systematically defined. Notably, dNK cells, which are abundant in early-pregnancy decidua, have been reported to show little or no expression of classical PGR [8]. Whether individual PAQR family members exhibit distinct cell-type-specific expression patterns across the maternal–fetal interface remains poorly characterized.

We therefore aimed to characterize the cell-type-specific expression of classical PGR and PAQR family members (PAQR5–PAQR9) at the human first-trimester maternal–fetal interface. We further examined the spatial relationships between selected PAQR expression patterns and the cell populations in which they were enriched, thereby characterizing the cellular and spatial architecture of classical and membrane progesterone receptor expression at the maternal–fetal interface.

## Methods

### Study design

We used a staged analytical framework to characterize classical *PGR* and *PAQR* family expression at the maternal–fetal interface. *PGR* and *PAQR5–PAQR9* expression was first mapped across cell populations in the first-trimester single-cell RNA-sequencing atlas of Vento-Tormo et al. [5]. Cell-type expression patterns were then evaluated in the Suryawanshi et al. first-trimester single-cell dataset after reference-based cell-type mapping [6]. Cell-type signatures were derived from the Vento-Tormo reference independently of *PGR* and *PAQR* expression, fixed before spatial analysis, and applied without reselection to Visium HD data. The prespecified spatial comparisons were *PAQR6* with a pan-dNK signature, *PAQR7* with villous cytotrophoblast (VCT) and syncytiotrophoblast (SCT) signatures, and *PAQR8* with Hofbauer cell (HB) and fetal fibroblast (fFB) signatures.

### Public single-cell datasets

For the discovery analysis, publicly available first-trimester placenta, decidua, and maternal blood single-cell data from Vento-Tormo et al. [5] (Nature 2018; ArrayExpress E-MTAB-6701 for the droplet-based dataset and E-MTAB-6678 for Smart-seq2) were used. A processed h5ad object was obtained from CELLxGENE. Raw counts were obtained from the AnnData raw/X layer after confirming that the values were non-negative and integer-like, and converted to a Seurat object while preserving the original metadata, author-provided cell-type labels, donor identifiers, tissue labels, assay labels, and the original embedding. No donor, tissue, or cell-type filtering was applied during object construction.

For cross-dataset evaluation, the 10x Genomics v2 dataset reported by Suryawanshi et al. [6] (Science Advances 2018; BioProject PRJNA492324) was obtained as a Human Cell Atlas Data Coordination Platform-generated loom file. The dataset contained two chorionic-villus preparations and one decidual preparation. Barcodes called as cells by EmptyDrops in the loom metadata were retained [9]. Cells with fewer than 200 detected genes, zero library size, or more than 25% mitochondrial transcripts were removed. Villous and decidual cells were processed separately for quality control.

### Vento-Tormo single-cell analysis

The principal discovery analysis used droplet-based 10x Genomics 3′ v2 data and the original author-provided cell-type annotations. Placental and decidual compartments were analyzed separately according to the tissue from which each cell was recovered; tissue origin itself was not used as a genotype-based assignment of fetal or maternal identity.

### Quantification and visualization of PGR and PAQR expression

For each cell, raw counts for *PGR* and *PAQR5–PAQR9* were normalized as ln[1 + (raw count / total cellular RNA count) × 10,000]. Within each cell-type group, detection frequency was defined as the percentage of cells with at least one raw count, and mean expression was calculated as the mean cell-wise log-normalized expression across all cells, including cells with zero counts. Dot size represents detection frequency, and dot color represents mean log-normalized expression. Open circles indicate groups in which no cell had detectable expression of the indicated transcript. Cell numbers were displayed beside each group. A common color scale was used within each dataset, whereas color scales were allowed to differ between datasets and were not used for direct quantitative comparison across studies.

### Cross-dataset single-cell reference mapping

To align the Suryawanshi dataset with the Vento-Tormo atlas without using the genes under study, a reference was constructed from the Vento-Tormo 10x placental and decidual cells. Original author-provided cell-type labels represented by at least 20 reference cells were eligible for mapping. To limit domination by abundant populations while retaining biological strata, each cell type was downsampled to a maximum of 400 cells, with representation balanced across tissue and donor. The reference was log-normalized with a scale factor of 10,000, and 3,000 variable features were selected. *PGR* and *PAQR5–PAQR9* were explicitly excluded from the reference feature set.

Reference anchors were identified using Seurat FindTransferAnchors with the reference principal component analysis (PCA) and the first 30 principal components, followed by TransferData [10]. A query cell was assigned a reference-mapped cell-type label only when the maximum prediction score was at least 0.50 and the difference between the highest and second-highest prediction scores was at least 0.10. Cells failing either criterion were designated unassigned and excluded from cell-type summaries. Only mapped groups containing at least 20 cells were plotted. The original query unique molecular identifier (UMI) counts were used to quantify *PGR* and *PAQR* expression as described above. T cells were retained as a pooled T-cell population because subtype-level separation was not sufficiently robust.

### Construction of PAQR-independent cell-type signatures

Cell-type signatures were constructed from the original Vento-Tormo droplet-based RNA count data before inspection of spatial correspondence. *PGR* and *PAQR5–PAQR9* were excluded from all candidate gene pools. Mitochondrial, ribosomal, globin, immunoglobulin, T-cell-receptor, HLA, KIR, sex-associated, and cell-cycle genes were also excluded. Counts were normalized using a scale factor of 10,000. Detection proportions and effect sizes were calculated, and one-versus-rest Wilcoxon rank-sum tests were performed using Seurat; adjusted *P* values were obtained using Seurat’s multiple-testing correction. Donor-level consistency was evaluated only after gene selection and was not used as a selection criterion. The final signature composition is summarized in Table 1, whereas comparator-level metrics for all prespecified contrasts are provided in Supplementary Table S1.

**Table 1.** PAQR-independent cell-type signatures used for spatial transcriptomic analysis. Cell-type signatures were derived from the Vento-Tormo et al. first-trimester single-cell RNA-sequencing dataset independently of PGR and PAQR expression and were fixed before spatial analysis.

| Cell-type signature | Signature composition | Selected genes | No. of genes | Prespecified spatial comparison |
| --- | --- | --- | --- | --- |
| VCT | De novo cell-type signature | SLC27A2, PARP1, SMAGP, ALDH7A1, ECI2, FAM3B, PDLIM1, BCAM | 8 | PAQR7–VCT |
| SCT | De novo cell-type signature | CYP19A1, LGALS16, GADD45G, LCMT1-AS2, ERVFRD-1, SERPINE1, LYPD3, MUC15 | 8 | PAQR7–SCT |
| HB | De novo cell-type signature | CD14, AIF1, C1QA, TYROBP, FOLR2, CCL3, C1QC, FCER1G | 8 | PAQR8–HB |
| fFB | De novo cell-type signature | DLK1, COL1A1, DCN, COL3A1, COL1A2, ACTA2, EGFL6, TAGLN | 8 | PAQR8–fFB |
| pan-dNK | 4 shared-core genes<br>+ 3 genes from each dNK subtype | Shared core: CMC1, CD7, IL2RB, KLRC1;<br>dNK1: SPINK2, CYP26A1, UBE2F; dNK2: XCL1, XCL2, ZNF683; dNK3: CCL5, ITM2C, CD160 | 13 | PAQR6–pan-dNK |

**Abbreviations:** VCT, villous cytotrophoblast; SCT, syncytiotrophoblast; HB, Hofbauer cell; fFB, fetal fibroblast; dNK, decidual natural killer cell.
**Note:** LCMT1-AS2 was retained in the prespecified eight-gene SCT signature but was unavailable in all three Visium HD feature matrices; therefore, spatial SCT scores were calculated using the remaining seven genes. Detailed comparator-level evidence for all selected signature genes is provided in Supplementary Table S1.

### Placental cell-type signatures

VCT, SCT, HB, and fFB signatures were derived de novo from the Vento-Tormo 10x placental cells. For each target population, candidate genes were required to be detected in at least 25% of target cells, to have an average log2 fold change (log2FC) of at least 0.50 and a difference in detection frequency between the target and comparator of at least 0.15 in every prespecified comparison, and to have an adjusted *P* value below 0.05 in the primary one-versus-rest Wilcoxon test. Prespecified comparisons were VCT versus all other placental cells, SCT, EVT, and pooled non-trophoblast cells; SCT versus all other placental cells, VCT, EVT, and pooled non-trophoblast cells; HB versus all other placental cells, pooled fFB1/fFB2, fetal endothelial cells, and pooled trophoblasts; and pooled fFB1/fFB2 versus all other placental cells, HB, fetal endothelial cells, and pooled trophoblasts.

To prevent the pooled fFB signature from being driven predominantly by either fibroblast subtype, genes were additionally required to be detected in at least 25% of both fFB1 and fFB2 cells and to show a log2FC of at least 0.25 for each subtype relative to non-fFB placental cells. Eligible genes were ranked first by the minimum log2FC across the required comparisons, followed by the minimum difference in detection frequency and the target-cell detection rate. The top eight genes were retained for each signature.

### Composite pan-dNK signature

The pan-dNK signature was designed to represent dNK1, dNK2, and dNK3 in a balanced manner. A shared core was identified genome-wide. Core genes were required to be detected in at least 25% of cells in each dNK subtype, to show a log2 fold change (log2FC) of at least 0.25 and a difference in detection frequency of at least 0.10 for each subtype versus pooled non-dNK decidual immune cells, and to show a pooled-dNK log2FC of at least 0.50, a pooled difference in detection frequency of at least 0.15, and an adjusted Wilcoxon *P* value below 0.05. Among eligible core genes, candidates were ranked by the minimum log2FC across the three individual subtype-versus-non-dNK comparisons, followed by the minimum difference in detection frequency, the minimum detection frequency across dNK1, dNK2, and dNK3, and gene symbol. The four highest-ranked core genes were retained.

For the subtype-specific components, candidate genes were restricted to those reported as positively differentially expressed for dNK1, dNK2, or dNK3 in Vento-Tormo et al. Supplementary Table 7 with an adjusted P value below 0.10. This published adjusted P value was used only for candidate prefiltering. For each subtype-specific component, the other two dNK subtypes were pooled to form a single within-dNK comparator. Each candidate was subsequently re-evaluated in the original 10x data in two separate contrasts: the target subtype versus the pooled other two dNK subtypes and the target subtype versus pooled non-dNK decidual immune cells. The latter comparator comprised the available T-cell, ILC3, decidual macrophage, dendritic-cell, monocyte, plasma-cell, granulocyte, and conventional NK-cell annotations in the decidual 10x dataset. Candidates were required to be detected in at least 25% of cells in the target subtype and to show a log2FC of at least 0.50 and a difference in detection frequency of at least 0.15 in each of the two required contrasts. Eligible genes were ranked by the smaller of the two log2FC values, followed by the smaller difference in detection frequency, target-subtype detection frequency, the published average logFC in Supplementary Table 7, and gene symbol. Three genes were retained for each subtype-specific component. Final subtype-arm selection was therefore based on detection frequency and effect-size criteria recalculated in the original 10x data. Proliferating dNK cells were excluded from signature construction and retained only as an audit population. The resulting pan-dNK signature comprised 13 genes: four shared-core genes and three genes representing each of the three dNK subtypes.

### Human tissue collection and Visium HD spatial transcriptomics

Maternal–fetal interface tissues were obtained from hysterectomy specimens from three pregnancies at gestational weeks (GW)11+0, GW11+6, and GW14+6. Hysterectomy was performed as part of surgical treatment for maternal ovarian or cervical cancer. None of the sampled placental or decidual tissues showed evidence of malignant involvement. The study was approved by the Institutional Review Boards of Kumamoto University (approval nos. 2566 and 2892) and Kyoto University (approval no. G0325). The use of archived tissue specimens was conducted under an opt-out consent procedure approved by the institutional review board.

Formalin-fixed, paraffin-embedded (FFPE) tissue sections were processed by Takara Bio Inc. using the 10x Genomics Visium HD Spatial Gene Expression workflow with Visium HD v1–FFPE chemistry, Visium HD Human Transcriptome 6.5-mm reagents, and the Dual Index Kit TS Set A. Following hematoxylin and eosin (H&E) staining and imaging, transcript-specific probe pairs were hybridized to tissue RNA and transferred to Visium HD slides using the Visium CytAssist. Libraries were sequenced on an Illumina NovaSeq 6000 using the NovaSeq 6000 S4 Reagent Kit v1.5 and NovaSeq Xp 4-Lane Kit v1.5 with paired-end sequencing comprising 43 bp for Read 1 and 50 bp for Read 2. FASTQ files were generated using spaceranger mkfastq v2.1.1. Spatial gene-expression data were processed using Space Ranger v3.0.1 with refdata-gex-GRCh38-2020-A and the Visium Human Transcriptome Probe Set v2.0 GRCh38-2020-A, comprising 18,085 included genes. All three tissue sections were processed using the same experimental and computational conditions. Filtered feature-barcode matrices from 8-µm square bins were used for downstream analyses.

### Visium HD processing and signature scoring

Three Visium HD sections were analyzed: GW11+0, GW11+6, and GW14+6, each from a separate case. Space Ranger filtered feature-barcode matrices from 8-µm square bins were imported using Seurat Read10X_h5. All filtered non-empty bins in each section were included in the analysis, and bins with zero total counts were removed.

Gene expression was calculated as ln[1 + (raw count / total bin UMI count) × 10,000]. The final cell-type signatures were fixed before spatial analysis and were not modified on the basis of *PAQR* spatial results (Table 1). *LCMT1-AS2*, one of the eight genes in the fixed SCT signature, was absent from all three Visium HD feature matrices; therefore, spatial SCT scores were calculated using the remaining seven genes. Within each section, each available fixed-signature gene was standardized across all filtered bins, and the composite signature score for each bin was calculated as the equal-weight mean of the gene-wise z scores. Genes unavailable in a given section or with zero variance were excluded from score calculation.

Signature-high bins were defined as bins at or above the section-specific 95th percentile of the composite signature score with at least one signature gene detected. Because ties at the cutoff could result in slightly more than 5% of bins being classified as signature-high, the realized fraction of signature-high bins was recorded. A *PAQR*-positive bin was defined as a bin containing at least one raw count for the target *PAQR* gene.

### Quantification of spatial correspondence

Spatial correspondence was quantified for the prespecified pairs *PAQR6*–pan-dNK, *PAQR7*–VCT, *PAQR7*–SCT, *PAQR8*–HB, and *PAQR8*–fFB. For each section, the numbers of *PAQR*-positive bins, signature-high bins, bins satisfying both definitions, and total filtered bins were recorded. The primary measure of spatial correspondence was the percentage of *PAQR*-positive bins that were also signature-high. Fold enrichment was calculated by dividing this percentage by the proportion of all filtered bins classified as signature-high. Thus, a fold enrichment greater than 1 indicates that signature-high bins were more frequent among *PAQR*-positive bins than in the section overall.

No bin-level hypothesis tests were performed because neighboring spatial bins are autocorrelated and do not represent independent biological replicates. Each tissue section was therefore treated as a descriptive biological replicate. Same-bin detection was interpreted as spatial correspondence rather than single-cell coexpression, because an 8-µm bin does not necessarily correspond to a single cell and may contain transcript signals from adjacent cells.

### Spatial visualization

For bivariate visualization, 8-µm square-bin data were displayed using Loupe Browser v8.1.2 (10x Genomics). Both axes represented the Loupe Browser LogNorm Feature Avg. The *PAQR* axis comprised the indicated single *PAQR* gene, whereas the cell-type-signature axis represented the average LogNorm expression across the available genes in the corresponding fixed signature. The visualization range for both axes was manually fixed from 0 to 4, with values above 4 displayed at maximum intensity. Increasing blue intensity indicated higher *PAQR* expression, increasing yellow intensity indicated higher signature-gene expression, and green indicated same-bin detection of signals on both axes. These Loupe Browser Feature Avg values were used solely for qualitative visualization and were not used to calculate the R-derived composite signature scores or to define signature-high bins.

## Results

### Cell-type-specific expression of classical PGR and PAQR family members

Analysis of the Vento-Tormo et al. single-cell RNA-sequencing dataset revealed distinct cell-type-specific expression patterns of classical *PGR* and membrane progestin receptor genes (*PAQR5–PAQR9*) across placental and decidual cell populations (Fig. 1A). *PGR* expression was predominantly detected in decidual stromal and perivascular populations, with detectable expression in approximately 12.6–35.1% of dS1–dS3 cells and 20.6–21.9% of dP1–dP2 cells, whereas expression was sparse or undetectable in most placental and decidual immune populations. In particular, *PGR* was detected in only 0–0.074% of dNK1–dNK3 cells.

**Figure 1.**
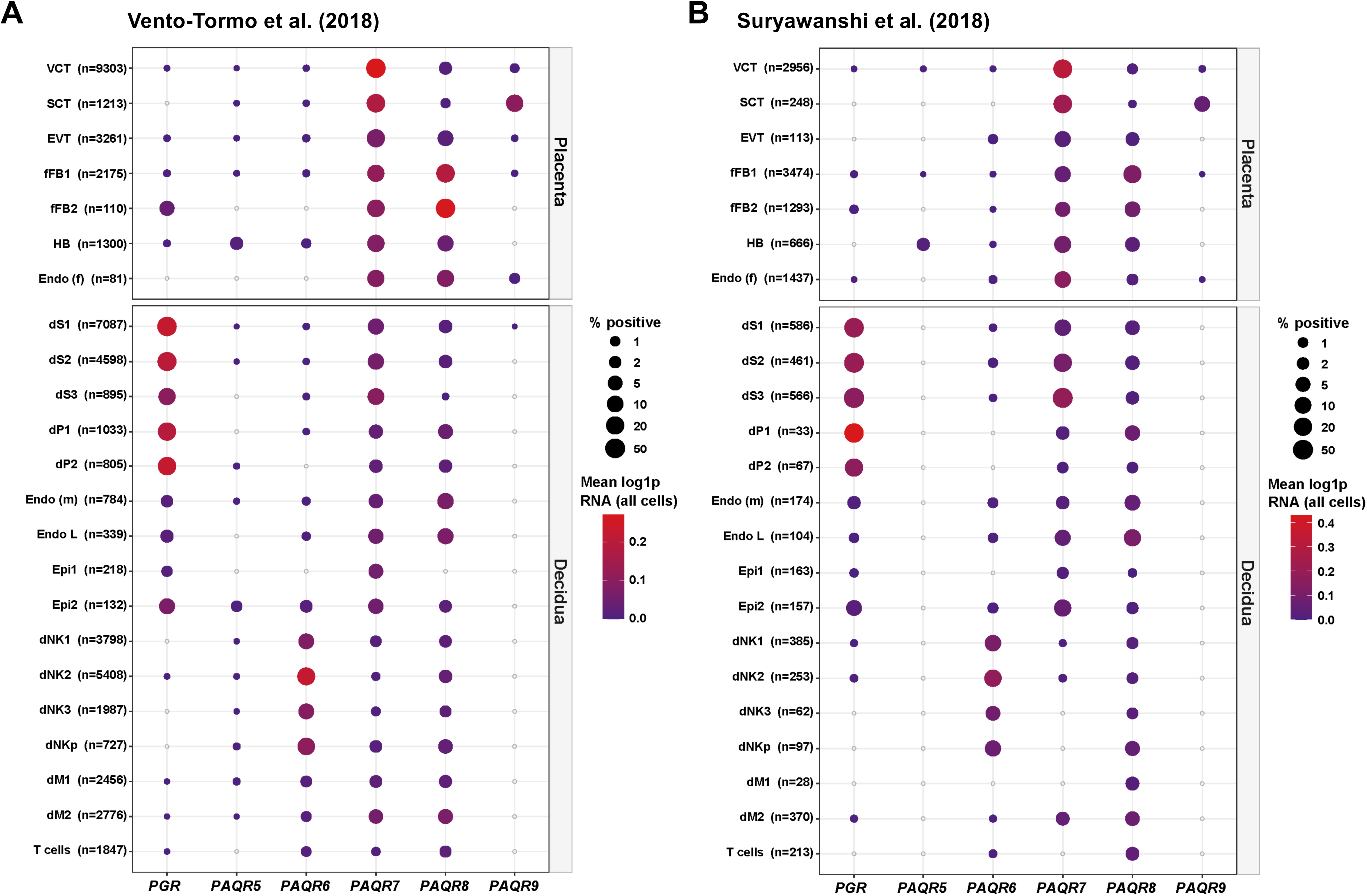
Cell-type-specific expression of classical and membrane progesterone receptors at the human maternal–fetal interface. **(A)** Expression of *PGR* and *PAQR5–PAQR9* in placental and decidual cell populations from the Vento-Tormo et al. (2018) scRNA-seq dataset. (B) Corresponding expression patterns in the Suryawanshi et al. (2018) scRNA-seq dataset, with cell-type labels transferred from the Vento-Tormo reference. Dot size represents the percentage of cells with detectable expression, and color represents mean log-normalized expression [ln(1 + counts per 10,000)] across all cells, including cells with zero counts. Open circles indicate no detectable expression. Cell numbers are shown in parentheses. Color scales are shown separately for each dataset and should not be used for direct quantitative comparison between datasets. Numbers denote transcriptionally defined subtypes within the indicated cell populations. VCT, villous cytotrophoblast; SCT, syncytiotrophoblast; EVT, extravillous trophoblast; fFB, fetal fibroblast; HB, Hofbauer cell; Endo (f), fetal endothelial cell; dS, decidual stromal cell; dP, decidual perivascular cell; Endo (m), maternal endothelial cell; Endo L, lymphatic endothelial cell; Epi, epithelial cell; dNK, decidual natural killer cell; dNKp, proliferating decidual natural killer cell; dM, decidual macrophage.

In contrast, individual *PAQR* family members exhibited distinct cell-type-specific expression patterns. *PAQR6* was enriched in dNK populations, with detectable expression in 7.1–17.5% of dNK1–dNK3 cells and 14.6% of dNKp cells. *PAQR7* was broadly expressed in placental trophoblast populations, including VCT (35.8%), SCT (22.8%), and EVT (19.3%). *PAQR8* showed a broader distribution, with prominent expression in fetal fibroblasts and Hofbauer cells as well as maternal decidual macrophages. *PAQR9* showed a more restricted pattern, with its strongest expression in SCT (13.5%). *PAQR5* was generally expressed at lower levels and showed no similarly prominent cell-type-specific enrichment.

The major expression patterns were also observed in the Suryawanshi et al. first-trimester single-cell RNA-sequencing dataset after cell-type labels were transferred from the Vento-Tormo reference (Fig. 1B). *PGR* was concentrated in decidual stromal and perivascular populations, *PAQR6* was enriched in dNK populations, *PAQR7* was prominent in trophoblasts, *PAQR8* was expressed across fetal stromal and macrophage populations, and *PAQR9* was enriched in SCT. These findings across two first-trimester single-cell datasets demonstrate a reproducible cell-type-specific organization of classical *PGR* and individual *PAQR* family members at the maternal–fetal interface.

### Spatial correspondence between PAQR expression and cell-type signatures

To examine the spatial relationships between PAQR expression and the cell populations identified by single-cell RNA sequencing, we analyzed three human maternal–fetal interface sections using Visium HD at 8-µm bin resolution. Spatial correspondence was assessed for prespecified receptor–cell type pairs based on the expression patterns observed in the single-cell datasets: *PAQR6* with pan-dNK, *PAQR7* with VCT and SCT, and *PAQR8* with HB and fFB signatures.

Representative spatial maps showed correspondence between *PAQR6* expression and the pan-dNK signature in decidua, *PAQR7* expression and VCT and SCT signatures in placental villi, and *PAQR8* expression and HB and fFB signatures within the villous stroma (Fig. 2A–E). These spatial relationships were observed across the three sections, although their magnitude varied among specimens.

**Figure 2.**
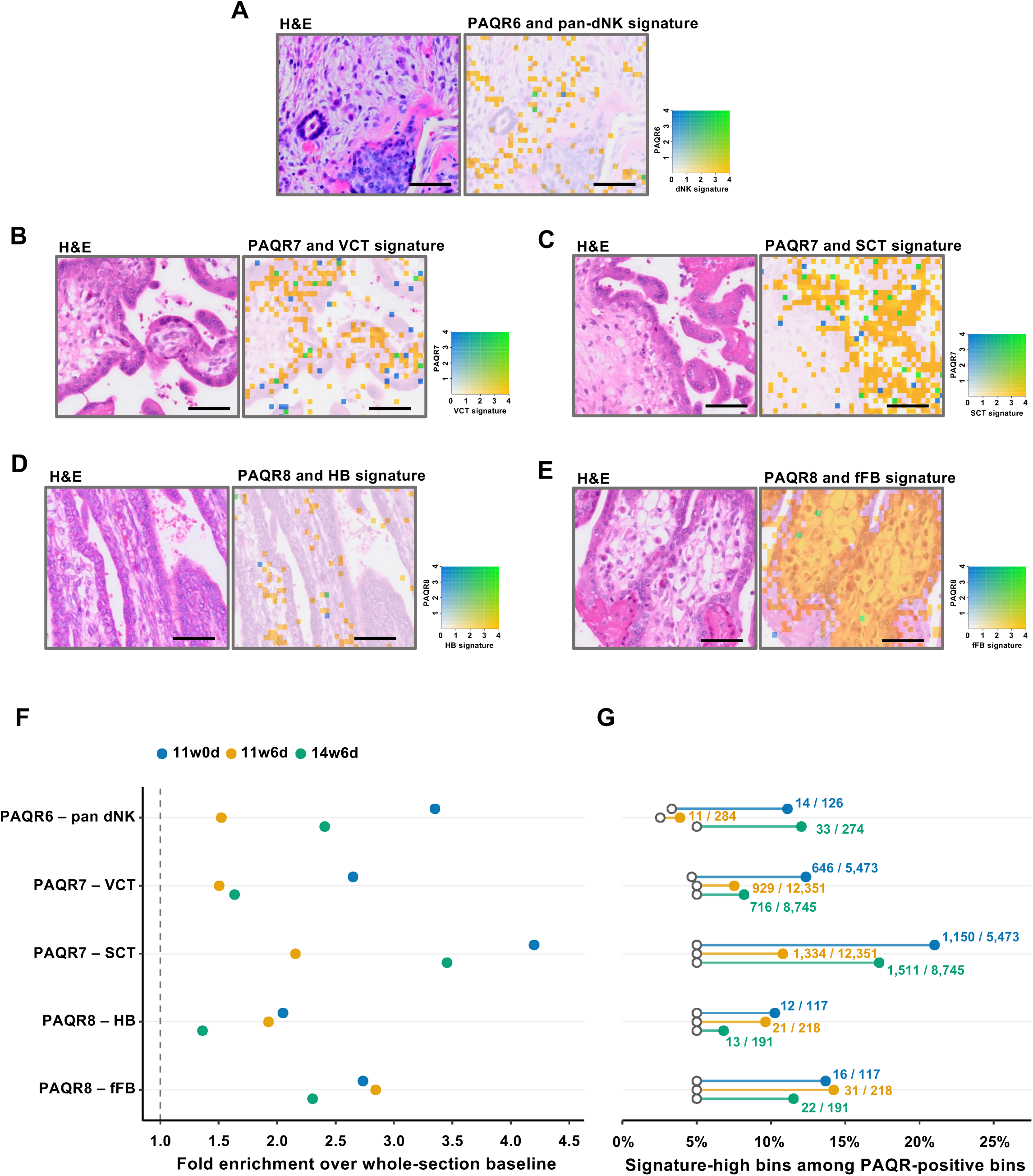
Spatial correspondence between PAQR expression and cell-type signatures at the human maternal–fetal interface. (A–E) Representative H&E images and corresponding bivariate Visium HD maps at 8-µm bin resolution, showing *PAQR6* and the pan-dNK signature in the GW14+6 section (A), *PAQR7* and the VCT (B) and SCT (C) signatures in the GW11+0 section, and *PAQR8* and the HB (D) and fFB (E) signatures in the GW11+6 section. Bivariate Loupe Browser maps show LogNorm expression within each 8-µm bin. The blue channel represents LogNorm expression of the indicated PAQR transcript, whereas the yellow channel represents the Feature Avg of LogNorm expression across the corresponding fixed cell-type signature genes. Both channels are displayed over a manually defined range of 0–4, with values above 4 shown at maximum intensity. Green indicates bins with signals on both channels. Scale bars, 50 µm. These Loupe Browser maps were used for qualitative visualization only and were not used to define signature-high bins in F and G. **(F)** Fold enrichment of signature-high bins among PAQR-positive bins relative to the proportion of signature-high bins among all filtered bins in the same section. The dashed vertical line indicates no enrichment (fold enrichment = 1). **(G)** Percentage of PAQR-positive bins classified as signature-high. Colored circles indicate the observed percentages, whereas open circles indicate the proportion of signature-high bins among all filtered bins in the same section; connecting lines show the difference between these values. Numbers indicate signature-high PAQR-positive bins/total PAQR-positive bins. Signature-high bins were defined as bins at or above the section-specific 95th percentile of composite signature scores, requiring detection of at least one signature gene, and PAQR-positive bins contained at least one UMI. Colors indicate the GW11+0, GW11+6, and GW14+6 sections. Each point represents one Visium HD section. Same-bin spatial correspondence does not establish single-cell coexpression.

Quantitative analysis demonstrated enrichment of signature-high bins among PAQR-positive bins relative to the whole-section prevalence of each cell-type signature (Fig. 2F). Enrichment was observed for all five prespecified receptor–cell type pairs in each section, with fold-enrichment values consistently exceeding 1. The strongest enrichment was observed for *PAQR7*–SCT and *PAQR6*–pan-dNK in some sections, whereas *PAQR8*–HB showed more modest enrichment.

Consistent with these findings, the proportion of PAQR-positive bins classified as signature-high exceeded the corresponding whole-section baseline for each receptor–cell type pair (Fig. 2G). Together, these spatial analyses support the cell-type-associated expression patterns identified by single-cell RNA sequencing and demonstrate spatial correspondence across the three tissue sections between *PAQR6* and dNK cells, *PAQR7* and trophoblast populations, and *PAQR8* and fetal stromal and macrophage populations at the maternal–fetal interface.

## Discussion

In this study, we delineated the cell-type-specific expression landscape of classical and membrane progesterone receptors at the human maternal–fetal interface during early pregnancy. Across two independent single-cell transcriptomic datasets, PGR expression was predominantly localized to decidual stromal and perivascular populations, whereas members of the PAQR family showed distinct and reproducible cellular distributions. PAQR6 was enriched in decidual natural killer cells, PAQR7 in trophoblast populations, PAQR8 in fetal stromal and macrophage compartments, and PAQR9 was most prominently expressed in syncytiotrophoblasts. Spatial transcriptomic analysis further supported the spatial correspondence of selected PAQR expression patterns with their associated cell-type signatures. Together, these findings reveal that progesterone receptor expression at the early maternal–fetal interface is highly cell-type specific and extends well beyond the distribution of the classical nuclear progesterone receptor.

Progesterone has long been recognized as an essential hormone for the establishment and maintenance of pregnancy; however, the physiological roles of membrane progesterone receptors during pregnancy remain largely unexplored. In addition to the well-established transcriptional actions mediated by the classical nuclear progesterone receptor, progesterone can elicit rapid, non-genomic responses through membrane-associated receptors, including members of the progestin and adipoQ receptor (PAQR) family [4]. Recent studies have begun to uncover previously unrecognized roles of these pathways in maternal adaptation and offspring development. In mice, maternal progesterone signaling through adipose mPRε/PAQR9 regulates maternal insulin resistance, thereby influencing nutrient availability to the developing embryo and postnatal metabolic homeostasis [11]. Moreover, mPRδ in the fetal submandibular gland senses maternal progesterone during pregnancy and, through DHA-mediated signaling, regulates salivary gland development and function as well as the establishment of the oral microbiota, thereby contributing to long-term oral homeostasis [12]. These findings illustrate how membrane progesterone receptors can translate the maternal endocrine environment into tissue-specific developmental and physiological responses. In this context, our finding that individual PAQR family members exhibit distinct cell-type-specific expression patterns at the early human maternal–fetal interface raises the possibility that membrane progesterone signaling similarly mediates specialized progesterone responses among trophoblast, immune, and stromal cell populations during early placentation.

The distinct cellular distribution of individual PAQR family members suggests that membrane progesterone signaling may be functionally compartmentalized at the maternal–fetal interface. A previous study using human gestational tissues reported abundant placental expression of mPRα/PAQR7 and PAQR9, with mPRα immunoreactivity strongly localized to syncytiotrophoblasts in term placental villi [7], supporting a role for membrane progesterone signaling in trophoblast biology. Our single-cell analyses extend these observations by resolving receptor expression across trophoblast subtypes, revealing prominent PAQR7 expression in trophoblast populations and a more selective enrichment of PAQR9 in syncytiotrophoblasts. This differential distribution raises the possibility that individual mPR subtypes contribute to distinct aspects of trophoblast differentiation or function. Particularly intriguing was the enrichment of PAQR6 in decidual natural killer (dNK) cells. dNK cells are a major immune population at the maternal–fetal interface and contribute to immune tolerance, vascular remodeling, and trophoblast–decidual interactions during early pregnancy [5,13]. Although progesterone is known to influence dNK-cell abundance and function, dNK cells have generally been considered to lack the classical nuclear progesterone receptor [8]. Progesterone effects on dNK cells have therefore been proposed to occur indirectly through decidual stromal cells or through alternative steroid receptors such as the glucocorticoid receptor [14]. The selective expression of PAQR6 in dNK cells identified here raises the possibility of an additional, direct pathway through which these cells may sense the progesterone-rich environment of early pregnancy. Finally, PAQR8 was preferentially expressed in fetal stromal and macrophage populations, including Hofbauer cells, suggesting that membrane progesterone responsiveness may also extend to the fetal mesenchymal and immune compartments of the developing placenta. Notably, the preferential expression of different PAQR subtypes in maternal dNK cells and fetal Hofbauer cells further suggests that progesterone sensing may be differentially configured between maternal and fetal immune compartments. Collectively, these cell-type-specific expression patterns support a model in which progesterone signaling at the maternal–fetal interface is not mediated by a single receptor pathway, but may instead be distributed among distinct cellular compartments through different progesterone receptor systems.

Several limitations of this study should be acknowledged. First, our analyses were based primarily on publicly available single-cell transcriptomic datasets and were therefore observational in nature; consequently, the present findings establish cellular expression patterns but do not demonstrate receptor-mediated progesterone signaling or its functional consequences. Second, transcript abundance does not necessarily reflect protein expression, subcellular localization, or receptor activity. Future studies combining protein-level validation with receptor-specific perturbation in relevant primary cells or experimental models will therefore be required to determine whether the identified PAQRs mediate direct cellular responses to progesterone. Third, our study focused on the early maternal–fetal interface, and whether these receptor expression patterns change with advancing gestation or under pregnancy complications remains unknown. Nevertheless, the reproducibility of the major expression patterns across independent single-cell datasets, together with the spatial correspondence observed in independent tissue sections, strengthens the robustness of the cellular map presented here and provides a foundation for future mechanistic studies of membrane progesterone signaling in human pregnancy.

In conclusion, this study provides a cell-type-resolved map of classical and membrane progesterone receptor expression at the human maternal–fetal interface during early pregnancy. The distinct distribution of PAQR family members across trophoblast, maternal immune, and fetal stromal and immune compartments provides a framework for understanding cell-type-specific progesterone signaling in human placentation.

## Author contributions

A.S. performed the bioinformatic analyses, interpreted the data, and wrote the manuscript; R.S., Y.I., M.K., M.Y., H.M., F.O., K.N., and H.O. interpreted the data; I.K. supervised the project and interpreted the data; E.K. interpreted the data and wrote the manuscript. E.K. had primary responsibility for the final content. All authors read and approved the final version of the manuscript.

## Data availability

The publicly available single-cell RNA-sequencing datasets analyzed in this study are available under the accession numbers described in the Methods. Data supporting the Visium HD analyses presented in this study are available from the corresponding author upon reasonable request. The complete Visium HD dataset is not publicly available at this time because it contains data related to ongoing, unpublished studies.

## References

1. Wetendorf M, DeMayo FJ. The progesterone receptor regulates implantation, decidualization, and glandular development via a complex paracrine signaling network. Mol Cell Endocrinol. 2012;357:108–118. doi:10.1016/j.mce.2011.10.028.

2. Motomura K, Miller D, Galaz J, Liu TN, Romero R, Gomez-Lopez N. The effects of progesterone on immune cellular function at the maternal-fetal interface and in maternal circulation. J Steroid Biochem Mol Biol. 2023;229:106254. doi:10.1016/j.jsbmb.2023.106254.

3. Large MJ, DeMayo FJ. The regulation of embryo implantation and endometrial decidualization by progesterone receptor signaling. Mol Cell Endocrinol. 2012;358:155–165. doi:10.1016/j.mce.2011.07.027.

4. Thomas P. Membrane progesterone receptors (mPRs, PAQRs): review of structural and signaling characteristics. Cells. 2022;11:1785. doi:10.3390/cells11111785.

5. Vento-Tormo R, Efremova M, Botting RA, et al. Single-cell reconstruction of the early maternal-fetal interface in humans. Nature. 2018;563:347–353. doi:10.1038/s41586-018-0698-6.

6. Suryawanshi H, Morozov P, Straus A, et al. A single-cell survey of the human first-trimester placenta and decidua. Sci Adv. 2018;4:eaau4788. doi:10.1126/sciadv.aau4788.

7. Fernandes MS, Pierron V, Michalovich D, et al. Regulated expression of putative membrane progestin receptor homologues in human endometrium and gestational tissues. J Endocrinol. 2005;187:89–101. doi:10.1677/joe.1.06242.

8. Henderson TA, Saunders PTK, Moffett-King A, Groome NP, Critchley HOD. Steroid receptor expression in uterine natural killer cells. J Clin Endocrinol Metab. 2003;88:440–449. doi:10.1210/jc.2002-021174.

9. Lun ATL, Riesenfeld S, Andrews T, et al. EmptyDrops: distinguishing cells from empty droplets in droplet-based single-cell RNA sequencing data. Genome Biol. 2019;20:63. doi:10.1186/s13059-019-1662-y.

10. Stuart T, Butler A, Hoffman P, et al. Comprehensive integration of single-cell data. Cell. 2019;177:1888–1902.e21. doi:10.1016/j.cell.2019.05.031.

11. Watanabe K, et al. Maternal progesterone and adipose mPRε in pregnancy regulate the embryonic nutritional state. Cell Rep. 2025;44:115433. doi:10.1016/j.celrep.2025.115433.

12. Yamano M, Miyamoto J, Sasahara D, et al. Maternal progesterone signaling establishes lifelong oral homeostasis. bioRxiv. 2026.08.10.743932. doi:10.64898/2026.08.10.743932.

13. Hanna J, Goldman-Wohl D, Hamani Y, et al. Decidual NK cells regulate key developmental processes at the human fetal-maternal interface. Nat Med. 2006;12:1065–1074. doi:10.1038/nm1452.

14. Guo W, Li P, Zhao G, Fan H, Hu Y, Hou Y. Glucocorticoid receptor mediates the effect of progesterone on uterine natural killer cells. Am J Reprod Immunol. 2012;67:463–473. doi:10.1111/j.1600-0897.2012.01114.x.

