## Supplementary_Table_S1 for "Cell-type-specific expression of classical and membrane progesterone receptors at the human maternal–fetal interface"

**Supplementary Table S1. Comparator-level evidence for PAQR-independent cell-type signatures**

One row is shown for each selected gene and required comparator. Blank adjusted P values indicate that no additional contrast-specific Wilcoxon test was performed; those contrasts were evaluated using prespecified detection and effect-size thresholds. For dNK subtype arms, the reported adjusted P value is the candidate-source value from Vento-Tormo et al. Supplementary Table 7.

| **Signature** | **Component** | **Rank** | **Gene** | **Contrast role** | **Target** | **Comparator** | **Target n** | **Comparator n** | **Target detection (%)** | **Comparator detection (%)** | **Δ detection (pp)** | **Mean norm. target** | **Mean norm. comparator** | **avg log2FC** | **Adjusted P value** | **P-value source** | **Min. required log2FC** | **Min. required Δ detection (pp)** | **Note** |
| --- | --- | --- | --- | --- | --- | --- | --- | --- | --- | --- | --- | --- | --- | --- | --- | --- | --- | --- | --- |
| VCT | VCT | 1 | SLC27A2 | Primary one-versus-rest contrast | VCT | All other selected fetal placental populations | 9303 | 8140 | 88.2 | 9.0 | 79.2 | 2.611 | 0.108 | 1.705 | <1 × 10^-300 | Primary one-versus-rest Wilcoxon test | 1.439 | 66.4 |  |
| VCT | VCT | 1 | SLC27A2 | Required lineage contrast | VCT | SCT | 9303 | 1213 | 88.2 | 21.8 | 66.4 | 2.611 | 0.331 | 1.439 |  | No additional contrast-specific Wilcoxon test; effect-size/detection threshold only | 1.439 | 66.4 |  |
| VCT | VCT | 1 | SLC27A2 | Required lineage contrast | VCT | EVT | 9303 | 3261 | 88.2 | 5.3 | 83.0 | 2.611 | 0.020 | 1.823 |  | No additional contrast-specific Wilcoxon test; effect-size/detection threshold only | 1.439 | 66.4 |  |
| VCT | VCT | 1 | SLC27A2 | Required lineage contrast | VCT | HB + fFB1 + fFB2 + Endo (f) | 9303 | 3666 | 88.2 | 8.1 | 80.1 | 2.611 | 0.112 | 1.700 |  | No additional contrast-specific Wilcoxon test; effect-size/detection threshold only | 1.439 | 66.4 |  |
| VCT | VCT | 2 | PARP1 | Primary one-versus-rest contrast | VCT | All other selected fetal placental populations | 9303 | 8140 | 97.3 | 59.3 | 38.0 | 4.450 | 0.850 | 1.559 | <1 × 10^-300 | Primary one-versus-rest Wilcoxon test | 1.392 | 30.1 |  |
| VCT | VCT | 2 | PARP1 | Required lineage contrast | VCT | SCT | 9303 | 1213 | 97.3 | 49.4 | 47.9 | 4.450 | 0.858 | 1.553 |  | No additional contrast-specific Wilcoxon test; effect-size/detection threshold only | 1.392 | 30.1 |  |
| VCT | VCT | 2 | PARP1 | Required lineage contrast | VCT | EVT | 9303 | 3261 | 97.3 | 67.2 | 30.1 | 4.450 | 0.594 | 1.774 |  | No additional contrast-specific Wilcoxon test; effect-size/detection threshold only | 1.392 | 30.1 |  |
| VCT | VCT | 2 | PARP1 | Required lineage contrast | VCT | HB + fFB1 + fFB2 + Endo (f) | 9303 | 3666 | 97.3 | 55.7 | 41.6 | 4.450 | 1.076 | 1.392 |  | No additional contrast-specific Wilcoxon test; effect-size/detection threshold only | 1.392 | 30.1 |  |
| VCT | VCT | 3 | SMAGP | Primary one-versus-rest contrast | VCT | All other selected fetal placental populations | 9303 | 8140 | 98.5 | 53.4 | 45.2 | 11.093 | 1.150 | 2.492 | <1 × 10^-300 | Primary one-versus-rest Wilcoxon test | 1.170 | 17.2 |  |
| VCT | VCT | 3 | SMAGP | Required lineage contrast | VCT | SCT | 9303 | 1213 | 98.5 | 81.4 | 17.2 | 11.093 | 4.374 | 1.170 |  | No additional contrast-specific Wilcoxon test; effect-size/detection threshold only | 1.170 | 17.2 |  |
| VCT | VCT | 3 | SMAGP | Required lineage contrast | VCT | EVT | 9303 | 3261 | 98.5 | 67.2 | 31.3 | 11.093 | 0.523 | 2.989 |  | No additional contrast-specific Wilcoxon test; effect-size/detection threshold only | 1.170 | 17.2 |  |
| VCT | VCT | 3 | SMAGP | Required lineage contrast | VCT | HB + fFB1 + fFB2 + Endo (f) | 9303 | 3666 | 98.5 | 31.8 | 66.8 | 11.093 | 0.641 | 2.881 |  | No additional contrast-specific Wilcoxon test; effect-size/detection threshold only | 1.170 | 17.2 |  |
| VCT | VCT | 4 | ALDH7A1 | Primary one-versus-rest contrast | VCT | All other selected fetal placental populations | 9303 | 8140 | 95.1 | 55.1 | 40.0 | 3.886 | 0.679 | 1.541 | <1 × 10^-300 | Primary one-versus-rest Wilcoxon test | 1.067 | 21.1 |  |
| VCT | VCT | 4 | ALDH7A1 | Required lineage contrast | VCT | SCT | 9303 | 1213 | 95.1 | 61.7 | 33.4 | 3.886 | 1.333 | 1.067 |  | No additional contrast-specific Wilcoxon test; effect-size/detection threshold only | 1.067 | 21.1 |  |
| VCT | VCT | 4 | ALDH7A1 | Required lineage contrast | VCT | EVT | 9303 | 3261 | 95.1 | 74.0 | 21.1 | 3.886 | 0.637 | 1.578 |  | No additional contrast-specific Wilcoxon test; effect-size/detection threshold only | 1.067 | 21.1 |  |
| VCT | VCT | 4 | ALDH7A1 | Required lineage contrast | VCT | HB + fFB1 + fFB2 + Endo (f) | 9303 | 3666 | 95.1 | 36.2 | 58.9 | 3.886 | 0.500 | 1.704 |  | No additional contrast-specific Wilcoxon test; effect-size/detection threshold only | 1.067 | 21.1 |  |
| VCT | VCT | 5 | ECI2 | Primary one-versus-rest contrast | VCT | All other selected fetal placental populations | 9303 | 8140 | 93.3 | 39.1 | 54.2 | 2.754 | 0.441 | 1.381 | <1 × 10^-300 | Primary one-versus-rest Wilcoxon test | 1.054 | 45.9 |  |
| VCT | VCT | 5 | ECI2 | Required lineage contrast | VCT | SCT | 9303 | 1213 | 93.3 | 47.4 | 45.9 | 2.754 | 0.808 | 1.054 |  | No additional contrast-specific Wilcoxon test; effect-size/detection threshold only | 1.054 | 45.9 |  |
| VCT | VCT | 5 | ECI2 | Required lineage contrast | VCT | EVT | 9303 | 3261 | 93.3 | 31.4 | 61.8 | 2.754 | 0.128 | 1.734 |  | No additional contrast-specific Wilcoxon test; effect-size/detection threshold only | 1.054 | 45.9 |  |
| VCT | VCT | 5 | ECI2 | Required lineage contrast | VCT | HB + fFB1 + fFB2 + Endo (f) | 9303 | 3666 | 93.3 | 43.1 | 50.2 | 2.754 | 0.599 | 1.231 |  | No additional contrast-specific Wilcoxon test; effect-size/detection threshold only | 1.054 | 45.9 |  |
| VCT | VCT | 6 | FAM3B | Primary one-versus-rest contrast | VCT | All other selected fetal placental populations | 9303 | 8140 | 90.6 | 17.7 | 72.9 | 2.934 | 0.231 | 1.676 | <1 × 10^-300 | Primary one-versus-rest Wilcoxon test | 1.025 | 39.3 |  |
| VCT | VCT | 6 | FAM3B | Required lineage contrast | VCT | SCT | 9303 | 1213 | 90.6 | 51.4 | 39.3 | 2.934 | 0.933 | 1.025 |  | No additional contrast-specific Wilcoxon test; effect-size/detection threshold only | 1.025 | 39.3 |  |
| VCT | VCT | 6 | FAM3B | Required lineage contrast | VCT | EVT | 9303 | 3261 | 90.6 | 10.0 | 80.7 | 2.934 | 0.039 | 1.920 |  | No additional contrast-specific Wilcoxon test; effect-size/detection threshold only | 1.025 | 39.3 |  |
| VCT | VCT | 6 | FAM3B | Required lineage contrast | VCT | HB + fFB1 + fFB2 + Endo (f) | 9303 | 3666 | 90.6 | 13.5 | 77.2 | 2.934 | 0.170 | 1.749 |  | No additional contrast-specific Wilcoxon test; effect-size/detection threshold only | 1.025 | 39.3 |  |
| VCT | VCT | 7 | PDLIM1 | Primary one-versus-rest contrast | VCT | All other selected fetal placental populations | 9303 | 8140 | 95.0 | 56.3 | 38.7 | 4.804 | 1.086 | 1.476 | <1 × 10^-300 | Primary one-versus-rest Wilcoxon test | 1.015 | 31.0 |  |
| VCT | VCT | 7 | PDLIM1 | Required lineage contrast | VCT | SCT | 9303 | 1213 | 95.0 | 53.9 | 41.1 | 4.804 | 1.872 | 1.015 |  | No additional contrast-specific Wilcoxon test; effect-size/detection threshold only | 1.015 | 31.0 |  |
| VCT | VCT | 7 | PDLIM1 | Required lineage contrast | VCT | EVT | 9303 | 3261 | 95.0 | 64.0 | 31.0 | 4.804 | 0.606 | 1.854 |  | No additional contrast-specific Wilcoxon test; effect-size/detection threshold only | 1.015 | 31.0 |  |
| VCT | VCT | 7 | PDLIM1 | Required lineage contrast | VCT | HB + fFB1 + fFB2 + Endo (f) | 9303 | 3666 | 95.0 | 50.2 | 44.8 | 4.804 | 1.253 | 1.365 |  | No additional contrast-specific Wilcoxon test; effect-size/detection threshold only | 1.015 | 31.0 |  |
| VCT | VCT | 8 | BCAM | Primary one-versus-rest contrast | VCT | All other selected fetal placental populations | 9303 | 8140 | 98.7 | 35.9 | 62.8 | 7.057 | 0.710 | 2.237 | <1 × 10^-300 | Primary one-versus-rest Wilcoxon test | 0.989 | 23.2 |  |
| VCT | VCT | 8 | BCAM | Required lineage contrast | VCT | SCT | 9303 | 1213 | 98.7 | 75.5 | 23.2 | 7.057 | 3.059 | 0.989 |  | No additional contrast-specific Wilcoxon test; effect-size/detection threshold only | 0.989 | 23.2 |  |
| VCT | VCT | 8 | BCAM | Required lineage contrast | VCT | EVT | 9303 | 3261 | 98.7 | 35.9 | 62.7 | 7.057 | 0.182 | 2.769 |  | No additional contrast-specific Wilcoxon test; effect-size/detection threshold only | 0.989 | 23.2 |  |
| VCT | VCT | 8 | BCAM | Required lineage contrast | VCT | HB + fFB1 + fFB2 + Endo (f) | 9303 | 3666 | 98.7 | 22.7 | 76.0 | 7.057 | 0.401 | 2.524 |  | No additional contrast-specific Wilcoxon test; effect-size/detection threshold only | 0.989 | 23.2 |  |
| SCT | SCT | 1 | CYP19A1 | Primary one-versus-rest contrast | SCT | All other selected fetal placental populations | 1213 | 16230 | 78.2 | 20.8 | 57.5 | 12.149 | 0.305 | 3.333 | <1 × 10^-300 | Primary one-versus-rest Wilcoxon test | 3.168 | 49.7 |  |
| SCT | SCT | 1 | CYP19A1 | Required lineage contrast | SCT | VCT | 1213 | 9303 | 78.2 | 28.5 | 49.7 | 12.149 | 0.463 | 3.168 |  | No additional contrast-specific Wilcoxon test; effect-size/detection threshold only | 3.168 | 49.7 |  |
| SCT | SCT | 1 | CYP19A1 | Required lineage contrast | SCT | EVT | 1213 | 3261 | 78.2 | 12.8 | 65.5 | 12.149 | 0.052 | 3.644 |  | No additional contrast-specific Wilcoxon test; effect-size/detection threshold only | 3.168 | 49.7 |  |
| SCT | SCT | 1 | CYP19A1 | Required lineage contrast | SCT | HB + fFB1 + fFB2 + Endo (f) | 1213 | 3666 | 78.2 | 8.3 | 69.9 | 12.149 | 0.129 | 3.542 |  | No additional contrast-specific Wilcoxon test; effect-size/detection threshold only | 3.168 | 49.7 |  |
| SCT | SCT | 2 | LGALS16 | Primary one-versus-rest contrast | SCT | All other selected fetal placental populations | 1213 | 16230 | 56.3 | 9.7 | 46.6 | 6.845 | 0.113 | 2.817 | <1 × 10^-300 | Primary one-versus-rest Wilcoxon test | 2.786 | 45.2 |  |
| SCT | SCT | 2 | LGALS16 | Required lineage contrast | SCT | VCT | 1213 | 9303 | 56.3 | 11.1 | 45.2 | 6.845 | 0.133 | 2.792 |  | No additional contrast-specific Wilcoxon test; effect-size/detection threshold only | 2.786 | 45.2 |  |
| SCT | SCT | 2 | LGALS16 | Required lineage contrast | SCT | EVT | 1213 | 3261 | 56.3 | 8.5 | 47.8 | 6.845 | 0.031 | 2.927 |  | No additional contrast-specific Wilcoxon test; effect-size/detection threshold only | 2.786 | 45.2 |  |
| SCT | SCT | 2 | LGALS16 | Required lineage contrast | SCT | HB + fFB1 + fFB2 + Endo (f) | 1213 | 3666 | 56.3 | 7.3 | 49.1 | 6.845 | 0.137 | 2.786 |  | No additional contrast-specific Wilcoxon test; effect-size/detection threshold only | 2.786 | 45.2 |  |
| SCT | SCT | 3 | GADD45G | Primary one-versus-rest contrast | SCT | All other selected fetal placental populations | 1213 | 16230 | 71.3 | 36.9 | 34.4 | 8.609 | 0.601 | 2.585 | 3.14e-272 | Primary one-versus-rest Wilcoxon test | 2.409 | 23.6 |  |
| SCT | SCT | 3 | GADD45G | Required lineage contrast | SCT | VCT | 1213 | 9303 | 71.3 | 47.7 | 23.6 | 8.609 | 0.810 | 2.409 |  | No additional contrast-specific Wilcoxon test; effect-size/detection threshold only | 2.409 | 23.6 |  |
| SCT | SCT | 3 | GADD45G | Required lineage contrast | SCT | EVT | 1213 | 3261 | 71.3 | 15.9 | 55.4 | 8.609 | 0.084 | 3.148 |  | No additional contrast-specific Wilcoxon test; effect-size/detection threshold only | 2.409 | 23.6 |  |
| SCT | SCT | 3 | GADD45G | Required lineage contrast | SCT | HB + fFB1 + fFB2 + Endo (f) | 1213 | 3666 | 71.3 | 28.2 | 43.1 | 8.609 | 0.531 | 2.650 |  | No additional contrast-specific Wilcoxon test; effect-size/detection threshold only | 2.409 | 23.6 |  |
| SCT | SCT | 4 | LCMT1-AS2 | Primary one-versus-rest contrast | SCT | All other selected fetal placental populations | 1213 | 16230 | 59.5 | 14.8 | 44.7 | 3.504 | 0.113 | 2.016 | <1 × 10^-300 | Primary one-versus-rest Wilcoxon test | 1.954 | 40.2 |  |
| SCT | SCT | 4 | LCMT1-AS2 | Required lineage contrast | SCT | VCT | 1213 | 9303 | 59.5 | 19.3 | 40.2 | 3.504 | 0.162 | 1.954 |  | No additional contrast-specific Wilcoxon test; effect-size/detection threshold only | 1.954 | 40.2 |  |
| SCT | SCT | 4 | LCMT1-AS2 | Required lineage contrast | SCT | EVT | 1213 | 3261 | 59.5 | 15.0 | 44.5 | 3.504 | 0.052 | 2.098 |  | No additional contrast-specific Wilcoxon test; effect-size/detection threshold only | 1.954 | 40.2 |  |
| SCT | SCT | 4 | LCMT1-AS2 | Required lineage contrast | SCT | HB + fFB1 + fFB2 + Endo (f) | 1213 | 3666 | 59.5 | 3.4 | 56.2 | 3.504 | 0.044 | 2.110 |  | No additional contrast-specific Wilcoxon test; effect-size/detection threshold only | 1.954 | 40.2 |  |
| SCT | SCT | 5 | ERVFRD-1 | Primary one-versus-rest contrast | SCT | All other selected fetal placental populations | 1213 | 16230 | 38.4 | 4.0 | 34.4 | 2.734 | 0.029 | 1.860 | <1 × 10^-300 | Primary one-versus-rest Wilcoxon test | 1.855 | 33.6 |  |
| SCT | SCT | 5 | ERVFRD-1 | Required lineage contrast | SCT | VCT | 1213 | 9303 | 38.4 | 4.2 | 34.2 | 2.734 | 0.032 | 1.855 |  | No additional contrast-specific Wilcoxon test; effect-size/detection threshold only | 1.855 | 33.6 |  |
| SCT | SCT | 5 | ERVFRD-1 | Required lineage contrast | SCT | EVT | 1213 | 3261 | 38.4 | 4.8 | 33.6 | 2.734 | 0.015 | 1.880 |  | No additional contrast-specific Wilcoxon test; effect-size/detection threshold only | 1.855 | 33.6 |  |
| SCT | SCT | 5 | ERVFRD-1 | Required lineage contrast | SCT | HB + fFB1 + fFB2 + Endo (f) | 1213 | 3666 | 38.4 | 2.7 | 35.7 | 2.734 | 0.032 | 1.855 |  | No additional contrast-specific Wilcoxon test; effect-size/detection threshold only | 1.855 | 33.6 |  |
| SCT | SCT | 6 | SERPINE1 | Primary one-versus-rest contrast | SCT | All other selected fetal placental populations | 1213 | 16230 | 77.0 | 26.7 | 50.3 | 5.910 | 0.545 | 2.161 | <1 × 10^-300 | Primary one-versus-rest Wilcoxon test | 1.850 | 40.8 |  |
| SCT | SCT | 6 | SERPINE1 | Required lineage contrast | SCT | VCT | 1213 | 9303 | 77.0 | 26.6 | 50.4 | 5.910 | 0.320 | 2.389 |  | No additional contrast-specific Wilcoxon test; effect-size/detection threshold only | 1.850 | 40.8 |  |
| SCT | SCT | 6 | SERPINE1 | Required lineage contrast | SCT | EVT | 1213 | 3261 | 77.0 | 36.2 | 40.8 | 5.910 | 0.917 | 1.850 |  | No additional contrast-specific Wilcoxon test; effect-size/detection threshold only | 1.850 | 40.8 |  |
| SCT | SCT | 6 | SERPINE1 | Required lineage contrast | SCT | HB + fFB1 + fFB2 + Endo (f) | 1213 | 3666 | 77.0 | 18.4 | 58.6 | 5.910 | 0.787 | 1.951 |  | No additional contrast-specific Wilcoxon test; effect-size/detection threshold only | 1.850 | 40.8 |  |
| SCT | SCT | 7 | LYPD3 | Primary one-versus-rest contrast | SCT | All other selected fetal placental populations | 1213 | 16230 | 88.4 | 47.5 | 40.9 | 5.531 | 0.640 | 1.994 | <1 × 10^-300 | Primary one-versus-rest Wilcoxon test | 1.839 | 17.0 |  |
| SCT | SCT | 7 | LYPD3 | Required lineage contrast | SCT | VCT | 1213 | 9303 | 88.4 | 53.8 | 34.5 | 5.531 | 0.825 | 1.839 |  | No additional contrast-specific Wilcoxon test; effect-size/detection threshold only | 1.839 | 17.0 |  |
| SCT | SCT | 7 | LYPD3 | Required lineage contrast | SCT | EVT | 1213 | 3261 | 88.4 | 71.4 | 17.0 | 5.531 | 0.663 | 1.974 |  | No additional contrast-specific Wilcoxon test; effect-size/detection threshold only | 1.839 | 17.0 |  |
| SCT | SCT | 7 | LYPD3 | Required lineage contrast | SCT | HB + fFB1 + fFB2 + Endo (f) | 1213 | 3666 | 88.4 | 10.2 | 78.2 | 5.531 | 0.150 | 2.505 |  | No additional contrast-specific Wilcoxon test; effect-size/detection threshold only | 1.839 | 17.0 |  |
| SCT | SCT | 8 | MUC15 | Primary one-versus-rest contrast | SCT | All other selected fetal placental populations | 1213 | 16230 | 71.5 | 33.8 | 37.7 | 4.703 | 0.425 | 2.001 | 1.62e-281 | Primary one-versus-rest Wilcoxon test | 1.775 | 25.0 |  |
| SCT | SCT | 8 | MUC15 | Required lineage contrast | SCT | VCT | 1213 | 9303 | 71.5 | 46.5 | 25.0 | 4.703 | 0.667 | 1.775 |  | No additional contrast-specific Wilcoxon test; effect-size/detection threshold only | 1.775 | 25.0 |  |
| SCT | SCT | 8 | MUC15 | Required lineage contrast | SCT | EVT | 1213 | 3261 | 71.5 | 30.4 | 41.1 | 4.703 | 0.144 | 2.317 |  | No additional contrast-specific Wilcoxon test; effect-size/detection threshold only | 1.775 | 25.0 |  |
| SCT | SCT | 8 | MUC15 | Required lineage contrast | SCT | HB + fFB1 + fFB2 + Endo (f) | 1213 | 3666 | 71.5 | 4.7 | 66.8 | 4.703 | 0.061 | 2.426 |  | No additional contrast-specific Wilcoxon test; effect-size/detection threshold only | 1.775 | 25.0 |  |
| HB | HB | 1 | CD14 | Primary one-versus-rest contrast | HB | All other selected fetal placental populations | 1300 | 16143 | 99.8 | 21.2 | 78.6 | 37.957 | 0.178 | 5.047 | <1 × 10^-300 | Primary one-versus-rest Wilcoxon test | 4.790 | 77.4 |  |
| HB | HB | 1 | CD14 | Required lineage contrast | HB | fFB1 + fFB2 | 1300 | 2285 | 99.8 | 14.1 | 85.7 | 37.957 | 0.285 | 4.922 |  | No additional contrast-specific Wilcoxon test; effect-size/detection threshold only | 4.790 | 77.4 |  |
| HB | HB | 1 | CD14 | Required lineage contrast | HB | Endo (f) | 1300 | 81 | 99.8 | 21.0 | 78.8 | 37.957 | 0.408 | 4.790 |  | No additional contrast-specific Wilcoxon test; effect-size/detection threshold only | 4.790 | 77.4 |  |
| HB | HB | 1 | CD14 | Required lineage contrast | HB | VCT + SCT + EVT | 1300 | 13777 | 99.8 | 22.3 | 77.4 | 37.957 | 0.159 | 5.071 |  | No additional contrast-specific Wilcoxon test; effect-size/detection threshold only | 4.790 | 77.4 |  |
| HB | HB | 2 | AIF1 | Primary one-versus-rest contrast | HB | All other selected fetal placental populations | 1300 | 16143 | 100.0 | 14.6 | 85.4 | 20.530 | 0.130 | 4.252 | <1 × 10^-300 | Primary one-versus-rest Wilcoxon test | 4.097 | 84.7 |  |
| HB | HB | 2 | AIF1 | Required lineage contrast | HB | fFB1 + fFB2 | 1300 | 2285 | 100.0 | 10.2 | 89.8 | 20.530 | 0.258 | 4.097 |  | No additional contrast-specific Wilcoxon test; effect-size/detection threshold only | 4.097 | 84.7 |  |
| HB | HB | 2 | AIF1 | Required lineage contrast | HB | Endo (f) | 1300 | 81 | 100.0 | 9.9 | 90.1 | 20.530 | 0.196 | 4.170 |  | No additional contrast-specific Wilcoxon test; effect-size/detection threshold only | 4.097 | 84.7 |  |
| HB | HB | 2 | AIF1 | Required lineage contrast | HB | VCT + SCT + EVT | 1300 | 13777 | 100.0 | 15.3 | 84.7 | 20.530 | 0.108 | 4.281 |  | No additional contrast-specific Wilcoxon test; effect-size/detection threshold only | 4.097 | 84.7 |  |
| HB | HB | 3 | C1QA | Primary one-versus-rest contrast | HB | All other selected fetal placental populations | 1300 | 16143 | 99.8 | 11.5 | 88.4 | 19.528 | 0.100 | 4.222 | <1 × 10^-300 | Primary one-versus-rest Wilcoxon test | 4.081 | 87.8 |  |
| HB | HB | 3 | C1QA | Required lineage contrast | HB | fFB1 + fFB2 | 1300 | 2285 | 99.8 | 8.1 | 91.8 | 19.528 | 0.213 | 4.081 |  | No additional contrast-specific Wilcoxon test; effect-size/detection threshold only | 4.081 | 87.8 |  |
| HB | HB | 3 | C1QA | Required lineage contrast | HB | Endo (f) | 1300 | 81 | 99.8 | 8.6 | 91.2 | 19.528 | 0.096 | 4.228 |  | No additional contrast-specific Wilcoxon test; effect-size/detection threshold only | 4.081 | 87.8 |  |
| HB | HB | 3 | C1QA | Required lineage contrast | HB | VCT + SCT + EVT | 1300 | 13777 | 99.8 | 12.0 | 87.8 | 19.528 | 0.081 | 4.247 |  | No additional contrast-specific Wilcoxon test; effect-size/detection threshold only | 4.081 | 87.8 |  |
| HB | HB | 4 | TYROBP | Primary one-versus-rest contrast | HB | All other selected fetal placental populations | 1300 | 16143 | 100.0 | 23.5 | 76.5 | 24.626 | 0.214 | 4.400 | <1 × 10^-300 | Primary one-versus-rest Wilcoxon test | 4.029 | 75.0 |  |
| HB | HB | 4 | TYROBP | Required lineage contrast | HB | fFB1 + fFB2 | 1300 | 2285 | 100.0 | 15.3 | 84.7 | 24.626 | 0.336 | 4.261 |  | No additional contrast-specific Wilcoxon test; effect-size/detection threshold only | 4.029 | 75.0 |  |
| HB | HB | 4 | TYROBP | Required lineage contrast | HB | Endo (f) | 1300 | 81 | 100.0 | 16.0 | 84.0 | 24.626 | 0.570 | 4.029 |  | No additional contrast-specific Wilcoxon test; effect-size/detection threshold only | 4.029 | 75.0 |  |
| HB | HB | 4 | TYROBP | Required lineage contrast | HB | VCT + SCT + EVT | 1300 | 13777 | 100.0 | 25.0 | 75.0 | 24.626 | 0.191 | 4.427 |  | No additional contrast-specific Wilcoxon test; effect-size/detection threshold only | 4.029 | 75.0 |  |
| HB | HB | 5 | FOLR2 | Primary one-versus-rest contrast | HB | All other selected fetal placental populations | 1300 | 16143 | 99.8 | 9.6 | 90.2 | 15.825 | 0.069 | 3.976 | <1 × 10^-300 | Primary one-versus-rest Wilcoxon test | 3.926 | 89.7 |  |
| HB | HB | 5 | FOLR2 | Required lineage contrast | HB | fFB1 + fFB2 | 1300 | 2285 | 99.8 | 6.7 | 93.1 | 15.825 | 0.107 | 3.926 |  | No additional contrast-specific Wilcoxon test; effect-size/detection threshold only | 3.926 | 89.7 |  |
| HB | HB | 5 | FOLR2 | Required lineage contrast | HB | Endo (f) | 1300 | 81 | 99.8 | 4.9 | 94.8 | 15.825 | 0.044 | 4.011 |  | No additional contrast-specific Wilcoxon test; effect-size/detection threshold only | 3.926 | 89.7 |  |
| HB | HB | 5 | FOLR2 | Required lineage contrast | HB | VCT + SCT + EVT | 1300 | 13777 | 99.8 | 10.1 | 89.7 | 15.825 | 0.063 | 3.985 |  | No additional contrast-specific Wilcoxon test; effect-size/detection threshold only | 3.926 | 89.7 |  |
| HB | HB | 6 | CCL3 | Primary one-versus-rest contrast | HB | All other selected fetal placental populations | 1300 | 16143 | 95.0 | 22.8 | 72.2 | 19.680 | 0.202 | 4.105 | <1 × 10^-300 | Primary one-versus-rest Wilcoxon test | 3.825 | 71.0 |  |
| HB | HB | 6 | CCL3 | Required lineage contrast | HB | fFB1 + fFB2 | 1300 | 2285 | 95.0 | 15.5 | 79.5 | 19.680 | 0.324 | 3.965 |  | No additional contrast-specific Wilcoxon test; effect-size/detection threshold only | 3.825 | 71.0 |  |
| HB | HB | 6 | CCL3 | Required lineage contrast | HB | Endo (f) | 1300 | 81 | 95.0 | 22.2 | 72.8 | 19.680 | 0.459 | 3.825 |  | No additional contrast-specific Wilcoxon test; effect-size/detection threshold only | 3.825 | 71.0 |  |
| HB | HB | 6 | CCL3 | Required lineage contrast | HB | VCT + SCT + EVT | 1300 | 13777 | 95.0 | 24.0 | 71.0 | 19.680 | 0.180 | 4.131 |  | No additional contrast-specific Wilcoxon test; effect-size/detection threshold only | 3.825 | 71.0 |  |
| HB | HB | 7 | C1QC | Primary one-versus-rest contrast | HB | All other selected fetal placental populations | 1300 | 16143 | 99.7 | 8.0 | 91.7 | 14.502 | 0.070 | 3.857 | <1 × 10^-300 | Primary one-versus-rest Wilcoxon test | 3.755 | 91.2 |  |
| HB | HB | 7 | C1QC | Required lineage contrast | HB | fFB1 + fFB2 | 1300 | 2285 | 99.7 | 5.3 | 94.4 | 14.502 | 0.149 | 3.755 |  | No additional contrast-specific Wilcoxon test; effect-size/detection threshold only | 3.755 | 91.2 |  |
| HB | HB | 7 | C1QC | Required lineage contrast | HB | Endo (f) | 1300 | 81 | 99.7 | 4.9 | 94.8 | 14.502 | 0.047 | 3.888 |  | No additional contrast-specific Wilcoxon test; effect-size/detection threshold only | 3.755 | 91.2 |  |
| HB | HB | 7 | C1QC | Required lineage contrast | HB | VCT + SCT + EVT | 1300 | 13777 | 99.7 | 8.5 | 91.2 | 14.502 | 0.057 | 3.875 |  | No additional contrast-specific Wilcoxon test; effect-size/detection threshold only | 3.755 | 91.2 |  |
| HB | HB | 8 | FCER1G | Primary one-versus-rest contrast | HB | All other selected fetal placental populations | 1300 | 16143 | 99.8 | 19.1 | 80.7 | 17.822 | 0.151 | 4.032 | <1 × 10^-300 | Primary one-versus-rest Wilcoxon test | 3.729 | 77.6 |  |
| HB | HB | 8 | FCER1G | Required lineage contrast | HB | fFB1 + fFB2 | 1300 | 2285 | 99.8 | 13.7 | 86.1 | 17.822 | 0.226 | 3.941 |  | No additional contrast-specific Wilcoxon test; effect-size/detection threshold only | 3.729 | 77.6 |  |
| HB | HB | 8 | FCER1G | Required lineage contrast | HB | Endo (f) | 1300 | 81 | 99.8 | 22.2 | 77.6 | 17.822 | 0.420 | 3.729 |  | No additional contrast-specific Wilcoxon test; effect-size/detection threshold only | 3.729 | 77.6 |  |
| HB | HB | 8 | FCER1G | Required lineage contrast | HB | VCT + SCT + EVT | 1300 | 13777 | 99.8 | 20.0 | 79.8 | 17.822 | 0.137 | 4.050 |  | No additional contrast-specific Wilcoxon test; effect-size/detection threshold only | 3.729 | 77.6 |  |
| fFB | fFB | 1 | DLK1 | Primary one-versus-rest contrast | fFB1 + fFB2 | All other selected fetal placental populations | 2285 | 15158 | 100.0 | 44.4 | 55.6 | 75.664 | 0.544 | 5.633 | <1 × 10^-300 | Primary one-versus-rest Wilcoxon test | 5.198 | 53.0 | Additional balance requirement: fFB1 detection = 100.0%; fFB2 detection = 100.0%; fFB1 log2FC vs non-fFB = 5.546; fFB2 log2FC vs non-fFB = 6.744 |
| fFB | fFB | 1 | DLK1 | Required lineage contrast | fFB1 + fFB2 | HB | 2285 | 1300 | 100.0 | 41.3 | 58.6 | 75.664 | 0.763 | 5.443 |  | No additional contrast-specific Wilcoxon test; effect-size/detection threshold only | 5.198 | 53.0 |  |
| fFB | fFB | 1 | DLK1 | Required lineage contrast | fFB1 + fFB2 | Endo (f) | 2285 | 81 | 100.0 | 46.9 | 53.0 | 75.664 | 1.088 | 5.198 |  | No additional contrast-specific Wilcoxon test; effect-size/detection threshold only | 5.198 | 53.0 |  |
| fFB | fFB | 1 | DLK1 | Required lineage contrast | fFB1 + fFB2 | VCT + SCT + EVT | 2285 | 13777 | 100.0 | 44.7 | 55.3 | 75.664 | 0.521 | 5.656 |  | No additional contrast-specific Wilcoxon test; effect-size/detection threshold only | 5.198 | 53.0 |  |
| fFB | fFB | 2 | COL1A1 | Primary one-versus-rest contrast | fFB1 + fFB2 | All other selected fetal placental populations | 2285 | 15158 | 1 | 36.0 | 64.0 | 55.562 | 0.448 | 5.288 | <1 × 10^-300 | Primary one-versus-rest Wilcoxon test | 4.150 | 62.9 | Additional balance requirement: fFB1 detection = 100.0%; fFB2 detection = 100.0%; fFB1 log2FC vs non-fFB = 5.327; fFB2 log2FC vs non-fFB = 4.150 |
| fFB | fFB | 2 | COL1A1 | Required lineage contrast | fFB1 + fFB2 | HB | 2285 | 1300 | 1 | 24.6 | 75.4 | 55.562 | 0.593 | 5.150 |  | No additional contrast-specific Wilcoxon test; effect-size/detection threshold only | 4.150 | 62.9 |  |
| fFB | fFB | 2 | COL1A1 | Required lineage contrast | fFB1 + fFB2 | Endo (f) | 2285 | 81 | 1 | 19.8 | 80.2 | 55.562 | 0.315 | 5.427 |  | No additional contrast-specific Wilcoxon test; effect-size/detection threshold only | 4.150 | 62.9 |  |
| fFB | fFB | 2 | COL1A1 | Required lineage contrast | fFB1 + fFB2 | VCT + SCT + EVT | 2285 | 13777 | 1 | 37.1 | 62.9 | 55.562 | 0.435 | 5.301 |  | No additional contrast-specific Wilcoxon test; effect-size/detection threshold only | 4.150 | 62.9 |  |
| fFB | fFB | 3 | DCN | Primary one-versus-rest contrast | fFB1 + fFB2 | All other selected fetal placental populations | 2285 | 15158 | 100.0 | 30.5 | 69.5 | 43.517 | 0.302 | 5.096 | <1 × 10^-300 | Primary one-versus-rest Wilcoxon test | 4.020 | 66.6 | Additional balance requirement: fFB1 detection = 100.0%; fFB2 detection = 100.0%; fFB1 log2FC vs non-fFB = 5.133; fFB2 log2FC vs non-fFB = 4.020 |
| fFB | fFB | 3 | DCN | Required lineage contrast | fFB1 + fFB2 | HB | 2285 | 1300 | 100.0 | 26.8 | 73.1 | 43.517 | 0.404 | 4.986 |  | No additional contrast-specific Wilcoxon test; effect-size/detection threshold only | 4.020 | 66.6 |  |
| fFB | fFB | 3 | DCN | Required lineage contrast | fFB1 + fFB2 | Endo (f) | 2285 | 81 | 100.0 | 33.3 | 66.6 | 43.517 | 0.436 | 4.954 |  | No additional contrast-specific Wilcoxon test; effect-size/detection threshold only | 4.020 | 66.6 |  |
| fFB | fFB | 3 | DCN | Required lineage contrast | fFB1 + fFB2 | VCT + SCT + EVT | 2285 | 13777 | 100.0 | 30.8 | 69.1 | 43.517 | 0.292 | 5.107 |  | No additional contrast-specific Wilcoxon test; effect-size/detection threshold only | 4.020 | 66.6 |  |
| fFB | fFB | 4 | COL3A1 | Primary one-versus-rest contrast | fFB1 + fFB2 | All other selected fetal placental populations | 2285 | 15158 | 100.0 | 44.8 | 55.2 | 65.230 | 0.588 | 5.382 | <1 × 10^-300 | Primary one-versus-rest Wilcoxon test | 3.924 | 18.5 | Additional balance requirement: fFB1 detection = 100.0%; fFB2 detection = 100.0%; fFB1 log2FC vs non-fFB = 5.408; fFB2 log2FC vs non-fFB = 4.746 |
| fFB | fFB | 4 | COL3A1 | Required lineage contrast | fFB1 + fFB2 | HB | 2285 | 1300 | 100.0 | 33.9 | 66.1 | 65.230 | 0.689 | 5.293 |  | No additional contrast-specific Wilcoxon test; effect-size/detection threshold only | 3.924 | 18.5 |  |
| fFB | fFB | 4 | COL3A1 | Required lineage contrast | fFB1 + fFB2 | Endo (f) | 2285 | 81 | 100.0 | 81.5 | 18.5 | 65.230 | 3.363 | 3.924 |  | No additional contrast-specific Wilcoxon test; effect-size/detection threshold only | 3.924 | 18.5 |  |
| fFB | fFB | 4 | COL3A1 | Required lineage contrast | fFB1 + fFB2 | VCT + SCT + EVT | 2285 | 13777 | 100.0 | 45.6 | 54.4 | 65.230 | 0.563 | 5.405 |  | No additional contrast-specific Wilcoxon test; effect-size/detection threshold only | 3.924 | 18.5 |  |
| fFB | fFB | 5 | COL1A2 | Primary one-versus-rest contrast | fFB1 + fFB2 | All other selected fetal placental populations | 2285 | 15158 | 99.9 | 33.6 | 66.3 | 33.398 | 0.397 | 4.622 | <1 × 10^-300 | Primary one-versus-rest Wilcoxon test | 3.911 | 36.4 | Additional balance requirement: fFB1 detection = 99.9%; fFB2 detection = 100.0%; fFB1 log2FC vs non-fFB = 4.649; fFB2 log2FC vs non-fFB = 3.959 |
| fFB | fFB | 5 | COL1A2 | Required lineage contrast | fFB1 + fFB2 | HB | 2285 | 1300 | 99.9 | 63.5 | 36.4 | 33.398 | 1.287 | 3.911 |  | No additional contrast-specific Wilcoxon test; effect-size/detection threshold only | 3.911 | 36.4 |  |
| fFB | fFB | 5 | COL1A2 | Required lineage contrast | fFB1 + fFB2 | Endo (f) | 2285 | 81 | 99.9 | 37.0 | 62.9 | 33.398 | 0.672 | 4.362 |  | No additional contrast-specific Wilcoxon test; effect-size/detection threshold only | 3.911 | 36.4 |  |
| fFB | fFB | 5 | COL1A2 | Required lineage contrast | fFB1 + fFB2 | VCT + SCT + EVT | 2285 | 13777 | 99.9 | 30.8 | 69.1 | 33.398 | 0.311 | 4.714 |  | No additional contrast-specific Wilcoxon test; effect-size/detection threshold only | 3.911 | 36.4 |  |
| fFB | fFB | 6 | ACTA2 | Primary one-versus-rest contrast | fFB1 + fFB2 | All other selected fetal placental populations | 2285 | 15158 | 78.6 | 18.6 | 60.1 | 18.478 | 0.158 | 4.072 | <1 × 10^-300 | Primary one-versus-rest Wilcoxon test | 3.732 | 44.1 | Additional balance requirement: fFB1 detection = 78.3%; fFB2 detection = 85.5%; fFB1 log2FC vs non-fFB = 4.035; fFB2 log2FC vs non-fFB = 4.662 |
| fFB | fFB | 6 | ACTA2 | Required lineage contrast | fFB1 + fFB2 | HB | 2285 | 1300 | 78.6 | 22.5 | 56.2 | 18.478 | 0.282 | 3.925 |  | No additional contrast-specific Wilcoxon test; effect-size/detection threshold only | 3.732 | 44.1 |  |
| fFB | fFB | 6 | ACTA2 | Required lineage contrast | fFB1 + fFB2 | Endo (f) | 2285 | 81 | 78.6 | 34.6 | 44.1 | 18.478 | 0.466 | 3.732 |  | No additional contrast-specific Wilcoxon test; effect-size/detection threshold only | 3.732 | 44.1 |  |
| fFB | fFB | 6 | ACTA2 | Required lineage contrast | fFB1 + fFB2 | VCT + SCT + EVT | 2285 | 13777 | 78.6 | 18.1 | 60.5 | 18.478 | 0.144 | 4.089 |  | No additional contrast-specific Wilcoxon test; effect-size/detection threshold only | 3.732 | 44.1 |  |
| fFB | fFB | 7 | EGFL6 | Primary one-versus-rest contrast | fFB1 + fFB2 | All other selected fetal placental populations | 2285 | 15158 | 99.8 | 12.0 | 87.8 | 25.417 | 0.121 | 4.558 | <1 × 10^-300 | Primary one-versus-rest Wilcoxon test | 3.618 | 72.7 | Additional balance requirement: fFB1 detection = 99.8%; fFB2 detection = 100.0%; fFB1 log2FC vs non-fFB = 4.505; fFB2 log2FC vs non-fFB = 5.335 |
| fFB | fFB | 7 | EGFL6 | Required lineage contrast | fFB1 + fFB2 | HB | 2285 | 1300 | 99.8 | 9.5 | 90.3 | 25.417 | 0.168 | 4.499 |  | No additional contrast-specific Wilcoxon test; effect-size/detection threshold only | 3.618 | 72.7 |  |
| fFB | fFB | 7 | EGFL6 | Required lineage contrast | fFB1 + fFB2 | Endo (f) | 2285 | 81 | 99.8 | 27.2 | 72.7 | 25.417 | 1.152 | 3.618 |  | No additional contrast-specific Wilcoxon test; effect-size/detection threshold only | 3.618 | 72.7 |  |
| fFB | fFB | 7 | EGFL6 | Required lineage contrast | fFB1 + fFB2 | VCT + SCT + EVT | 2285 | 13777 | 99.8 | 12.1 | 87.7 | 25.417 | 0.111 | 4.572 |  | No additional contrast-specific Wilcoxon test; effect-size/detection threshold only | 3.618 | 72.7 |  |
| fFB | fFB | 8 | TAGLN | Primary one-versus-rest contrast | fFB1 + fFB2 | All other selected fetal placental populations | 2285 | 15158 | 77.8 | 23.2 | 54.6 | 15.499 | 0.383 | 3.576 | <1 × 10^-300 | Primary one-versus-rest Wilcoxon test | 3.564 | 50.7 | Additional balance requirement: fFB1 detection = 77.5%; fFB2 detection = 84.5%; fFB1 log2FC vs non-fFB = 3.564; fFB2 log2FC vs non-fFB = 3.793 |
| fFB | fFB | 8 | TAGLN | Required lineage contrast | fFB1 + fFB2 | HB | 2285 | 1300 | 77.8 | 19.5 | 58.3 | 15.499 | 0.255 | 3.717 |  | No additional contrast-specific Wilcoxon test; effect-size/detection threshold only | 3.564 | 50.7 |  |
| fFB | fFB | 8 | TAGLN | Required lineage contrast | fFB1 + fFB2 | Endo (f) | 2285 | 81 | 77.8 | 27.2 | 50.7 | 15.499 | 0.373 | 3.587 |  | No additional contrast-specific Wilcoxon test; effect-size/detection threshold only | 3.564 | 50.7 |  |
| fFB | fFB | 8 | TAGLN | Required lineage contrast | fFB1 + fFB2 | VCT + SCT + EVT | 2285 | 13777 | 77.8 | 23.6 | 54.3 | 15.499 | 0.395 | 3.564 |  | No additional contrast-specific Wilcoxon test; effect-size/detection threshold only | 3.564 | 50.7 |  |
| pan-dNK | core | 1 | CMC1 | Primary pooled contrast | Pooled dNK1 + dNK2 + dNK3 | Pooled non-dNK decidual immune cells | 11193 | 8047 | 93.5 | 43.5 | 50.0 |  |  | 2.713 | <1 × 10^-300 | Pooled dNK versus pooled non-dNK Wilcoxon test | 2.557 | 48.0 | Shared-core gene |
| pan-dNK | core | 1 | CMC1 | Required subtype contrast | dNK1 | Pooled non-dNK decidual immune cells | 3798 | 8047 | 95.6 | 43.5 | 52.1 |  |  | 2.798 |  | No additional contrast-specific Wilcoxon test; effect-size/detection threshold only | 2.557 | 48.0 | Shared-core gene |
| pan-dNK | core | 1 | CMC1 | Required subtype contrast | dNK2 | Pooled non-dNK decidual immune cells | 5408 | 8047 | 92.7 | 43.5 | 49.3 |  |  | 2.557 |  | No additional contrast-specific Wilcoxon test; effect-size/detection threshold only | 2.557 | 48.0 | Shared-core gene |
| pan-dNK | core | 1 | CMC1 | Required subtype contrast | dNK3 | Pooled non-dNK decidual immune cells | 1987 | 8047 | 91.5 | 43.5 | 48.0 |  |  | 2.932 |  | No additional contrast-specific Wilcoxon test; effect-size/detection threshold only | 2.557 | 48.0 | Shared-core gene |
| pan-dNK | core | 2 | CD7 | Primary pooled contrast | Pooled dNK1 + dNK2 + dNK3 | Pooled non-dNK decidual immune cells | 11193 | 8047 | 92.8 | 48.0 | 44.7 |  |  | 2.658 | <1 × 10^-300 | Pooled dNK versus pooled non-dNK Wilcoxon test | 2.181 | 39.9 | Shared-core gene |
| pan-dNK | core | 2 | CD7 | Required subtype contrast | dNK1 | Pooled non-dNK decidual immune cells | 3798 | 8047 | 87.9 | 48.0 | 39.9 |  |  | 2.181 |  | No additional contrast-specific Wilcoxon test; effect-size/detection threshold only | 2.181 | 39.9 | Shared-core gene |
| pan-dNK | core | 2 | CD7 | Required subtype contrast | dNK2 | Pooled non-dNK decidual immune cells | 5408 | 8047 | 95.3 | 48.0 | 47.3 |  |  | 2.987 |  | No additional contrast-specific Wilcoxon test; effect-size/detection threshold only | 2.181 | 39.9 | Shared-core gene |
| pan-dNK | core | 2 | CD7 | Required subtype contrast | dNK3 | Pooled non-dNK decidual immune cells | 1987 | 8047 | 95.0 | 48.0 | 47.0 |  |  | 2.412 |  | No additional contrast-specific Wilcoxon test; effect-size/detection threshold only | 2.181 | 39.9 | Shared-core gene |
| pan-dNK | core | 3 | IL2RB | Primary pooled contrast | Pooled dNK1 + dNK2 + dNK3 | Pooled non-dNK decidual immune cells | 11193 | 8047 | 92.8 | 31.6 | 61.2 |  |  | 2.451 | <1 × 10^-300 | Pooled dNK versus pooled non-dNK Wilcoxon test | 2.148 | 59.1 | Shared-core gene |
| pan-dNK | core | 3 | IL2RB | Required subtype contrast | dNK1 | Pooled non-dNK decidual immune cells | 3798 | 8047 | 96.6 | 31.6 | 65.0 |  |  | 2.843 |  | No additional contrast-specific Wilcoxon test; effect-size/detection threshold only | 2.148 | 59.1 | Shared-core gene |
| pan-dNK | core | 3 | IL2RB | Required subtype contrast | dNK2 | Pooled non-dNK decidual immune cells | 5408 | 8047 | 90.7 | 31.6 | 59.1 |  |  | 2.216 |  | No additional contrast-specific Wilcoxon test; effect-size/detection threshold only | 2.148 | 59.1 | Shared-core gene |
| pan-dNK | core | 3 | IL2RB | Required subtype contrast | dNK3 | Pooled non-dNK decidual immune cells | 1987 | 8047 | 91.2 | 31.6 | 59.6 |  |  | 2.148 |  | No additional contrast-specific Wilcoxon test; effect-size/detection threshold only | 2.148 | 59.1 | Shared-core gene |
| pan-dNK | core | 4 | KLRC1 | Primary pooled contrast | Pooled dNK1 + dNK2 + dNK3 | Pooled non-dNK decidual immune cells | 11193 | 8047 | 92.6 | 30.7 | 61.9 |  |  | 2.969 | <1 × 10^-300 | Pooled dNK versus pooled non-dNK Wilcoxon test | 2.024 | 45.3 | Shared-core gene |
| pan-dNK | core | 4 | KLRC1 | Required subtype contrast | dNK1 | Pooled non-dNK decidual immune cells | 3798 | 8047 | 96.8 | 30.7 | 66.1 |  |  | 3.067 |  | No additional contrast-specific Wilcoxon test; effect-size/detection threshold only | 2.024 | 45.3 | Shared-core gene |
| pan-dNK | core | 4 | KLRC1 | Required subtype contrast | dNK2 | Pooled non-dNK decidual immune cells | 5408 | 8047 | 95.8 | 30.7 | 65.0 |  |  | 3.141 |  | No additional contrast-specific Wilcoxon test; effect-size/detection threshold only | 2.024 | 45.3 | Shared-core gene |
| pan-dNK | core | 4 | KLRC1 | Required subtype contrast | dNK3 | Pooled non-dNK decidual immune cells | 1987 | 8047 | 76.0 | 30.7 | 45.3 |  |  | 2.024 |  | No additional contrast-specific Wilcoxon test; effect-size/detection threshold only | 2.024 | 45.3 | Shared-core gene |
| pan-dNK | dNK1 | 5 | SPINK2 | Required within-dNK contrast | dNK1 | Pooled other two dNK subtypes | 3798 | 7395 | 78.7 | 35.1 | 43.6 |  |  | 2.112 | <1 × 10^-300 | Nature 2018 Supplementary Table 7 positive-DE adjusted P (candidate-source) | 2.112 | 43.6 | Subtype-arm rank 1; re-evaluated in original 10x data |
| pan-dNK | dNK1 | 5 | SPINK2 | Required non-dNK contrast | dNK1 | Pooled non-dNK decidual immune cells | 3798 | 8047 | 78.7 | 21.2 | 57.4 |  |  | 3.053 |  | No additional contrast-specific Wilcoxon test; effect-size/detection threshold only | 2.112 | 43.6 | Subtype-arm rank 1; re-evaluated in original 10x data |
| pan-dNK | dNK1 | 6 | CYP26A1 | Required within-dNK contrast | dNK1 | Pooled other two dNK subtypes | 3798 | 7395 | 45.3 | 2.5 | 42.8 |  |  | 1.601 | <1 × 10^-300 | Nature 2018 Supplementary Table 7 positive-DE adjusted P (candidate-source) | 1.590 | 42.2 | Subtype-arm rank 2; re-evaluated in original 10x data |
| pan-dNK | dNK1 | 6 | CYP26A1 | Required non-dNK contrast | dNK1 | Pooled non-dNK decidual immune cells | 3798 | 8047 | 45.3 | 3.1 | 42.2 |  |  | 1.590 |  | No additional contrast-specific Wilcoxon test; effect-size/detection threshold only | 1.590 | 42.2 | Subtype-arm rank 2; re-evaluated in original 10x data |
| pan-dNK | dNK1 | 7 | UBE2F | Required within-dNK contrast | dNK1 | Pooled other two dNK subtypes | 3798 | 7395 | 72.7 | 33.1 | 39.6 |  |  | 1.537 | <1 × 10^-300 | Nature 2018 Supplementary Table 7 positive-DE adjusted P (candidate-source) | 1.537 | 39.6 | Subtype-arm rank 3; re-evaluated in original 10x data |
| pan-dNK | dNK1 | 7 | UBE2F | Required non-dNK contrast | dNK1 | Pooled non-dNK decidual immune cells | 3798 | 8047 | 72.7 | 27.0 | 45.8 |  |  | 1.805 |  | No additional contrast-specific Wilcoxon test; effect-size/detection threshold only | 1.537 | 39.6 | Subtype-arm rank 3; re-evaluated in original 10x data |
| pan-dNK | dNK2 | 8 | XCL1 | Required within-dNK contrast | dNK2 | Pooled other two dNK subtypes | 5408 | 5785 | 96.8 | 65.7 | 31.1 |  |  | 1.654 | <1 × 10^-300 | Nature 2018 Supplementary Table 7 positive-DE adjusted P (candidate-source) | 1.654 | 31.1 | Subtype-arm rank 1; re-evaluated in original 10x data |
| pan-dNK | dNK2 | 8 | XCL1 | Required non-dNK contrast | dNK2 | Pooled non-dNK decidual immune cells | 5408 | 8047 | 96.8 | 37.0 | 59.8 |  |  | 3.528 |  | No additional contrast-specific Wilcoxon test; effect-size/detection threshold only | 1.654 | 31.1 | Subtype-arm rank 1; re-evaluated in original 10x data |
| pan-dNK | dNK2 | 9 | XCL2 | Required within-dNK contrast | dNK2 | Pooled other two dNK subtypes | 5408 | 5785 | 97.5 | 78.2 | 19.3 |  |  | 1.585 | <1 × 10^-300 | Nature 2018 Supplementary Table 7 positive-DE adjusted P (candidate-source) | 1.585 | 19.3 | Subtype-arm rank 2; re-evaluated in original 10x data |
| pan-dNK | dNK2 | 9 | XCL2 | Required non-dNK contrast | dNK2 | Pooled non-dNK decidual immune cells | 5408 | 8047 | 97.5 | 41.2 | 56.3 |  |  | 3.619 |  | No additional contrast-specific Wilcoxon test; effect-size/detection threshold only | 1.585 | 19.3 | Subtype-arm rank 2; re-evaluated in original 10x data |
| pan-dNK | dNK2 | 10 | ZNF683 | Required within-dNK contrast | dNK2 | Pooled other two dNK subtypes | 5408 | 5785 | 39.5 | 12.3 | 27.3 |  |  | 0.986 | 3e-253 | Nature 2018 Supplementary Table 7 positive-DE adjusted P (candidate-source) | 0.986 | 27.3 | Subtype-arm rank 3; re-evaluated in original 10x data |
| pan-dNK | dNK2 | 10 | ZNF683 | Required non-dNK contrast | dNK2 | Pooled non-dNK decidual immune cells | 5408 | 8047 | 39.5 | 7.7 | 31.8 |  |  | 1.407 |  | No additional contrast-specific Wilcoxon test; effect-size/detection threshold only | 0.986 | 27.3 | Subtype-arm rank 3; re-evaluated in original 10x data |
| pan-dNK | dNK3 | 11 | CCL5 | Required within-dNK contrast | dNK3 | Pooled other two dNK subtypes | 1987 | 9206 | 94.0 | 59.3 | 34.7 |  |  | 2.598 | <1 × 10^-300 | Nature 2018 Supplementary Table 7 positive-DE adjusted P (candidate-source) | 2.598 | 34.7 | Subtype-arm rank 1; re-evaluated in original 10x data |
| pan-dNK | dNK3 | 11 | CCL5 | Required non-dNK contrast | dNK3 | Pooled non-dNK decidual immune cells | 1987 | 8047 | 94.0 | 40.1 | 53.9 |  |  | 2.796 |  | No additional contrast-specific Wilcoxon test; effect-size/detection threshold only | 2.598 | 34.7 | Subtype-arm rank 1; re-evaluated in original 10x data |
| pan-dNK | dNK3 | 12 | ITM2C | Required within-dNK contrast | dNK3 | Pooled other two dNK subtypes | 1987 | 9206 | 35.5 | 5.0 | 30.5 |  |  | 1.919 | <1 × 10^-300 | Nature 2018 Supplementary Table 7 positive-DE adjusted P (candidate-source) | 1.564 | 22.3 | Subtype-arm rank 2; re-evaluated in original 10x data |
| pan-dNK | dNK3 | 12 | ITM2C | Required non-dNK contrast | dNK3 | Pooled non-dNK decidual immune cells | 1987 | 8047 | 35.5 | 13.2 | 22.3 |  |  | 1.564 |  | No additional contrast-specific Wilcoxon test; effect-size/detection threshold only | 1.564 | 22.3 | Subtype-arm rank 2; re-evaluated in original 10x data |
| pan-dNK | dNK3 | 13 | CD160 | Required within-dNK contrast | dNK3 | Pooled other two dNK subtypes | 1987 | 9206 | 31.0 | 1.2 | 29.8 |  |  | 1.498 | <1 × 10^-300 | Nature 2018 Supplementary Table 7 positive-DE adjusted P (candidate-source) | 1.393 | 28.2 | Subtype-arm rank 3; re-evaluated in original 10x data |
| pan-dNK | dNK3 | 13 | CD160 | Required non-dNK contrast | dNK3 | Pooled non-dNK decidual immune cells | 1987 | 8047 | 31.0 | 2.8 | 28.2 |  |  | 1.393 |  | No additional contrast-specific Wilcoxon test; effect-size/detection threshold only | 1.393 | 28.2 | Subtype-arm rank 3; re-evaluated in original 10x data |
